# AHLF: ad hoc learning of peptide fragmentation from mass spectra enables an interpretable detection of phosphorylated and cross-linked peptides

**DOI:** 10.1101/2020.05.19.101345

**Authors:** Tom Altenburg, Sven Giese, Shengbo Wang, Thilo Muth, Bernhard Y. Renard

## Abstract

Mass spectrometry-based proteomics provides a holistic snapshot of the entire protein set of a living cell on a molecular level. Currently, only a few deep learning approaches that involve peptide fragmentation spectra, which represent partial sequence information of proteins, exist. Commonly, these approaches lack the ability to characterize less studied or even unknown patterns in spectra because of their use of explicit domain knowledge. To elevate unrestricted learning from spectra, we introduce AHLF, a deep learning model that is end-to-end trained on 19.2 million spectra from multiple phosphoproteomic data sets. AHLF is interpretable and we show that peak-level feature importances and pairwise interactions between peaks are in line with corresponding peptide fragments. We demonstrate our approach by detecting post-translational modifications, specifically protein phosphorylation based on only the fragmentation spectrum without a database search. AHLF increases the area under the receiver operating characteristic curve (AUC) by an average of 9.4% on recent phosphoproteomic data compared to the current-state-of-the-art on this task. To show the broad applicability of AHLF we use transfer learning to also detect cross-linked peptides, as used in protein structure analysis, with an AUC of up to 94%. We expect our approach to directly apply to cell signaling and structural biology which use phosphoproteomic and cross-linking data, but in principal any mass spectrometry based study can benefit from an interpretable, end-to-end trained model like AHLF.

**Availability:** https://gitlab.com/dacs-hpi/ahlf

## Introduction

Publicly available mass spectrometry (MS)-based proteomics data has grown exponentially in terms of the number of data sets and amount of data [1]. This is due to high-throughput proteomic studies generating a vast pool of fragmentation spectra. Each spectrum contains characteristic peak patterns which appear due to the fragmentation of a given peptide. These peptides are used to study the proteins contained in the biological sample. It might seem obvious to apply deep learning to solve various problems on this wealth of data, but the application of deep learning directly to fragmentation spectra has not yet sparked in the community. Instead, the MS wet lab workflow is usually followed by a conventional database search [2]. In a search, each acquired mass spectrum is scored against a list of candidate peptides from in-silico digested proteins. For each peptide candidate, a theoretical spectrum is constructed and compared to the acquired spectrum [3].

But the identification of spectra remains challenging as proteins are often either mutated or carry post-translational modifications (PTMs). The latter are essential for various biological processes and protein phosphorylation is an important PTM, regulating protein function and facilitating cellular signaling [4, 5]. Various sophisticated algorithms exist in order to cope with the challenges that arise from PTMs and mutations but still require protein databases [6, 7, 8, 9].

There are attempts to make predictions, e.g. detecting a PTM, based on only the spectrum itself and are therefore independent of a database. However, current approaches are based on engineered features and classical machine learning [10, 11]. A fragmentation spectrum can contain PTM-specific patterns (e.g. relations between peaks in a spectrum) that coexist with fragments resulting from the plain peptide sequence [12]. These patterns can even appear in an equivariant manner, i.e. they can pinpoint the position of a PTM within the peptide sequence but their presence alone can reveal the PTM itself [13]. Most importantly, the detection of a PTM can be separated from the sequence retrieval and a deep learning approach would account for the variety and complexity of PTM-specific patterns.

Aside from this, there is a plethora of open challenges in MS-based proteomics, including the detection of PTMs [12], prediction of phosphosite localization scores [13], detection of cross-linked peptides [14], characterization of the dark matter in proteomics by assigning spectrum-identifyability scores [15], augmenting features for post-search re-scoring [16], identification of biomarkers [17], detection of anomalies including non-proteinogenic amino acids [18] simply to name a few. These may be solved by a deep learning approach, when the underlying model is able to gather biochemically relevant reasoning from a large pool of spectra. As a proof of concept, we tackle two of the above mentioned issues, including the detection of spectra from peptides with PTMs and spectra from cross-linked peptides.

Regarding biological data, deep learning approaches on imaging or sequential data are very successfully published in a high frequency. We argue this is mainly due to the straight-forward applicability of findings from computer vision and natural language processing to medical image [19] or genomic data [20]. Interestingly, there are at least two applications of deep learning models that are well-received by the proteomics community: i) fragment intensity prediction and ii) *de novo* sequencing. Yet, the models that are used for both applications are predominantly informed by peptide sequences rather than spectra alone. A recurrent neural network applied to a peptide sequence predicts fragment intensities as implemented in pDeep [21] and Prosit [22]. Similarly, a recurrent neural network facilitates *de novo* sequencing by guiding a dynamic programming approach with a learned heuristic of peptide sequence patterns in case of DeepNovo [23, 24]. The latter also relies on a convolutional neural network that extracts features from the mass spectrum. But it is unable to characterize basic fragmentation patterns on its own, i.e. the various ion-types of amino acids are built-in and not trainable parameters.

Connected to this, is the lack of interpretability for these models because the built-in features prevent any further interpretation altogether. Predefined features may reflect the experts experience but restrict the flexibility of the model and effectively reduce interpretablility [25]. Hence, a generic deep learning model without engineered features would be beneficial for interpretation as it would primarily allow an unbiased view on the structure inherent in the data itself.

To the best of our knowledge, there have been no attempts of directly presenting fragmentation mass spectra to a deep learning model and ad hoc learn fragmentation patterns. Here, ad hoc learning means purposely abstracting essential fragmentation patterns in a data-driven but task-specific manner. For example, a PTM-detection would require the learning of modification-specific features, which we can further investigate after a model has been trained.

Therefore, we propose AHLF (‘ad hoc learning fragmentation’), an end-to-end trained deep neural network that learns from fragmentation spectra in order to perform versatile prediction tasks. To perform these challenging prediction tasks, AHLF features: i) a framework enabling efficient training on large numbers of fragmentation spectra, ii) the learning of long-range peak associations through dilated convolutions iii) true end-to-end training on spectra due to not including domain knowledge iv) interpretation of AHLF investigating whether biochemically relevant patterns are recognized.

We evaluate our approach on previously published phosphoproteomic data. This includes data sets from more than one hundred public repositories [4] and additionally validate these results on recently published data [26, 27, 28, 29, 30] and in case of cross-linking data we use previously published data to evaluate our transfer learning approach [31, 32, 33].

We interpret our model by comparing peptide fragments from ground truth peptide identifications against feature importance values for AHLF, which we calculated for each peak per spectrum individually on a collection of spectra. This is enabled by applying the SHAP framework [34] to predictions from AHLF. These feature importance values explain AHLF but also indicate potential discrepancies within the ground truth peptide identifications. Additionally, we show that AHLF recognizes biochemically relevant fragmentation patterns. In this case, we do not require annotations from identified peptides beforehand. We achieve this by computing pairwise interactions, using Path Explain [35], between any two peaks per spectrum and subsequently identify relevant delta masses between respective peak-pairs.

Finally, we demonstrate the broad scope of our approach by applying AHLF to a distinct task, the detection of cross-linked peptides (AHLFx). Cross-linking is used to study the structure of single-proteins, multiprotein complexes or protein-protein interactions [36]. Detecting spectra from cross-linked peptides is challenging and different from PTM-detection because in this case two peptides including the cross-linker molecule are present in the same spectrum and need to be detected. To the extent of our knowledge, this is the first instance where an immediate detection of spectra from potential cross-linked peptides is performed prior to database search. Additionally, this transfer learning approach demonstrates the applicability of AHLF for a situation where fewer labeled spectra are available for training.

## Results

### Spectrum representation of AHLF: allows deep learning on fragmentation mass spectra

A key challenge for deep learning on proteomics data is the representation of spectra. AHLF exploits the sparsity of fragmentation spectra to derive a memory-efficient representation that accounts for exact peak locations (Fig. 1).

**Figure 1:**
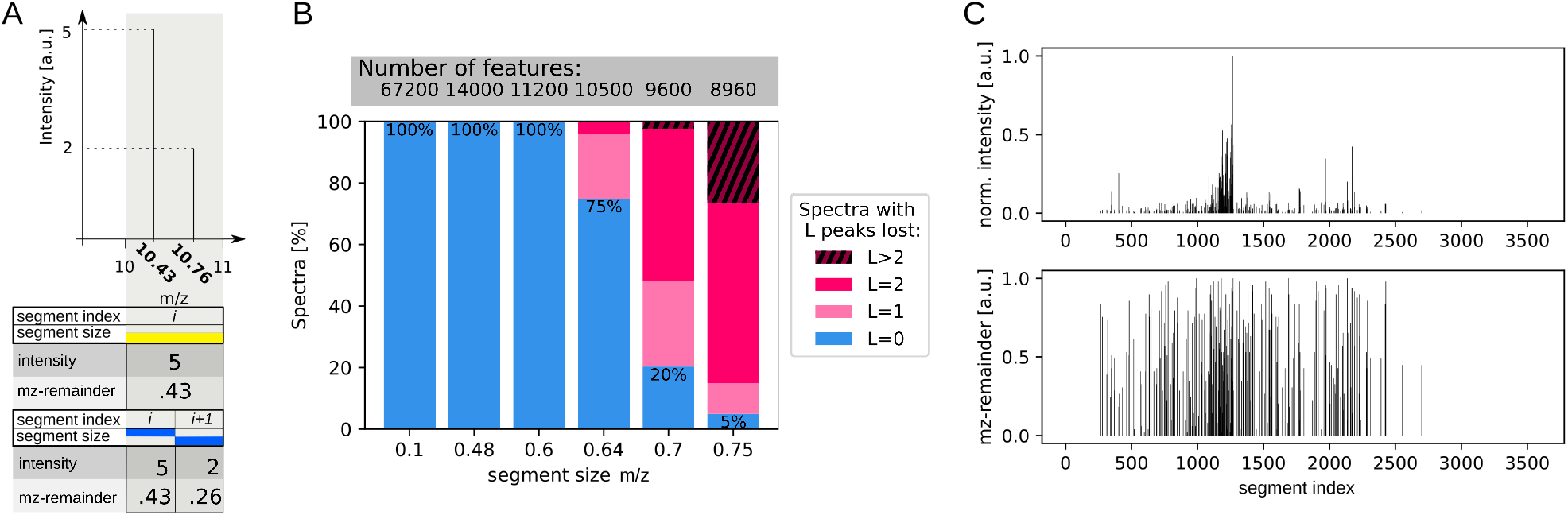
Two-vector spectrum representation is memory-efficient but keeps exact m/z locations. **A:** Example spectrum of two hypothetical peaks (top). Feature representation as two-vector spectrum (bottom): for a too large and thus lossy (1 m/z) segment size (yellow) and a smaller, loss-less (0.5 m/z) segment size (blue). **B:** Trade-off between number of features (top, with a windows-size 100-3560 m/z) and loss of peaks (y-axis), depending on the chosen segment size (x-axis). Note, the striped area at the top (L>2) of the stacked bars for the two widest segment sizes reflects spectra with more than two peaks lost. **C:** Fragmentation mass spectrum represented in two-vectors: intensity (top) and mz-remainder (bottom). The two feature vectors have 7200 features in total. From the mz-remainder the original m/z values can be fully recovered.

Mass spectrometers are highly accurate in performing measurements of mass over charge (m/z). For example in Fig. 1B, the fragmentation spectra from the Lung data set of PXD012174, have been acquired with a resolution of 7500. This means two peaks at 200 m/z can be separated if they differ in at least 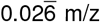. A naive conversion of such a spectrum into a dense feature vector, using this error and given window size (range of acquired m/z values) between 100 m/z and 3560 m/z, would result in a vector containing 129.750 features. In theory, such a naive representation is compatible with convolutions or recurrent operations. In practice, to reach acceptable training times, it is necessary to keep the number of features low while retaining most or even all information.

In each spectrum only a characteristic subset of all possible m/z is observed and those have non-zero intensities assigned. We exploit this sparsity here, for each peak (tuple of m/z and intensity) in a given spectrum we calculate the mz-remainder *m/z*′ according to (equation 1): *m/z* = *i · s* + *m/z*′, where *i* is an integer that specifies the segment index with fixed segment size *s* (Fig. 1A). Furthermore, *m/z*′ < *s* such that the tuple (*i*, *m/z*′) describes the location of a peak at *m/z*. If more than one peak is falling into the same segment, only the highest peak is kept (Fig. 1A and pseudocode supplementary algorithm 1). This yields two feature vectors, one containing intensities and the other the mz-remainder *m/z*′, the exact position within the respective segment, Fig. 1C. Note the recoverability of exact peak locations (original m/z values) of kept peaks from our two-vector spectrum is guaranteed by equation 1.

The segment size controls two important aspects of the two-vector spectrum representation: (i) the risk to lose peaks, (ii) the total number of features. The trade-off between these two aspects is shown in Fig. 1B. In fact, for more than 1.2 million spectra studied in Fig. 1B, not a single spectrum lost a peak when we use segments smaller or equal to 0.6 m/z. For this survey in Fig. 1B we used all spectra (including unidentified ones) from the Lung data set of PXD012174 [4], which in-turn was used for cross-validation below.

Based on Fig. 1B we selected a segment size of 0.5 m/z. Additionally, we chose a window size of 100-1900 m/z as peaks appear less frequent for larger m/z (this is common domain knowledge and therefore not explicitly shown here). The resulting two-vector spectrum has 7200 features in total, which is a reduction of number of features by factor 18 compared to the naive representation above. Hereinafter, all spectra are represented as a two-vector spectrum which is a tensor of size (3600,2). For the kept peaks, the original m/z values can be fully recovered and thus the information of exact peak locations can be used by AHLF, which we empirically demonstrate below.

### AHLF promotes learning of long-range peak associations via dilated convolutions

In general, peaks of peptide fragments are scattered over an entire spectrum. Hence, we designed our deep learning model in order to promote the learning of associations between any peaks while respecting their location within a spectrum Fig. 2. Therefore, we use convolutional layers because they preserve the location of a feature as the outputs from a convolution are equivariant with respect to their inputs. Subsequently, the higher layers of our deep neural network can make use of the presence and the location of peaks. To be exact, we use convolutions with gaps, commonly called dilated convolutions. These have a larger receptive field than vanilla convolutions but both have the same number of trainable parameters. In particular, our network has a receptive field that spans the entire feature vector. This is not only due to the use of dilated convolutions alone, but rather the receptive field grows exponentially with the number of layers Fig. 2. This is facilitated by a dilation rate (size of the gap; steps with zeros between trainable weights of convolutional filter) that increases alongside additional layers. Ultimately, due to parameter-sharing properties of convolutions, the total number of trainable weights is low compared to a fully connected network. Overall, the model architecture allows AHLF to use the entire two-vector spectrum as-is.

**Figure 2.**
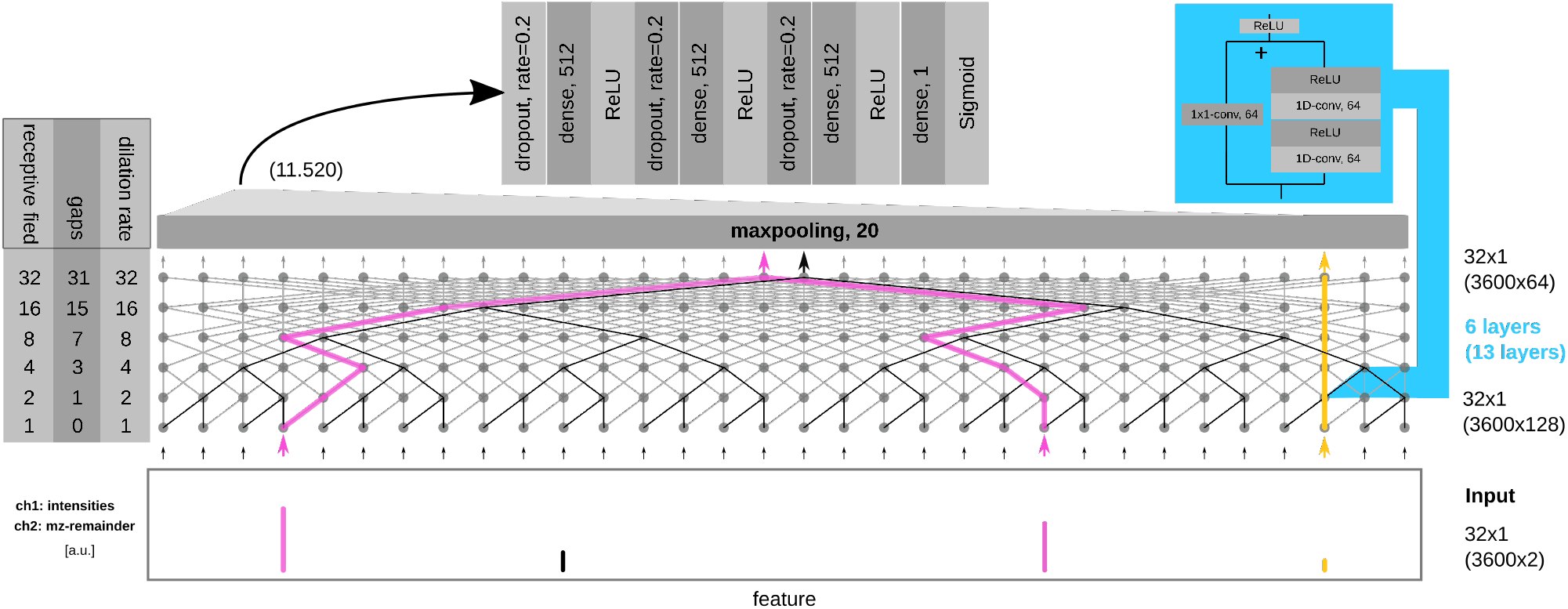
Illustration of long-range association learning via dilated convolutions in the deep network AHLF. Hyperparameters are summarized in table S1.1. The receptive field grows exponentially with additional layers according to specific dilation rate (box left). A possible information-flow (black solid lines): a single output (top) can receive information from any input feature (bottom) and indicates that the receptive field spans the entire feature vector. Nested stacks of two stacked convolutions and a skip-connection further increase non-linearity (blue, right). Only selected parts of the actual network are illustrated here. In parentheses (on the right) the actual tensor size and numbers of filters are compared against what is illustrated (without parentheses) here. We exemplify learnable associations: (i) between a specific pair of ions (pink path) or (ii) simply passing inputs through skip-connections (yellow path), other associations are possible to be learned (not illustrated here). An example of a hypothetical feature vector (bottom) with four peaks is shown, the two-vector representation is presented in two channels (ch1+ch2) to the model AHLF.

AHLF has 6,705,985 trainable model parameters of which 256,832 are part of the dilated convolutions. A summary of hyperparameters can be found in the supplementary table S1.1. Details for model training, hyperparameters selection including a training history of our final models can be found in the Methods section and supplementary material Fig. S2.2. AHLF computes prediction scores for 1000 spectra in 1.085 seconds, on a machine with a Nvidia V100 16GB GPU and an Intel Xeon processor with 14 CPU cores.

### AHLF robustly detects phosphorylations of peptides based on their fragmentation spectra

Here, we exemplify our approach on a specific use-case by applying AHLF to phosphoproteomic data and detect spectra of phosphorylated peptides. We compare our approach against PhoStar [11], which is a random forest model that includes carefully generated phospho-specific features. In contrast, AHLF was applied to the data in a plug-and-play manner as AHLF did not require any domain knowledge beforehand. Rather AHLF has to come up with domain specific features on its own, which we further investigate by interpreting AHLF later in this work.

To demonstrate the ability of AHLF to detect spectra of phosphorylated peptides by evaluating the performance of AHLF on 19.2 million labeled spectra from 101 cell/tissue-types available from 112 individual PRIDE repositories (PXD012174 [4]). The data set is roughly balanced and includes 10.5 million phosphorylated and 8.7 million unphosphorylated peptide-spectrum-matches (PSMs). We performed a four-fold cross validation yielding four independently trained deep learning models AHLF-*α*, −*β*, −*γ* and −*δ* with their respective hold-out fold **a,b,c** and **d**, see supplementary tables S1.2 and S1.3. The following results were computed by applying each model to its respective hold-out fold. For convenience we refer to AHLF in any case. We computed binary prediction scores and computed balanced accuracy (Bacc), F1-score and ROC-AUC for each of the 101 individual data sets. As an aggregation over these sets we show mean, median and variance in table 1 (detailed metrics for individual data sets are given in supplementary Tables S1.5).

**Table 1:** Median, mean and variance for ROC-AUC, F1-score and Bacc over the individual data sets.

|  | median ROC-AUC | median F1 | median Bacc |
| --- | --- | --- | --- |
| AHLF | **88.48** | **83.32** | **76.80** |
| PhoStar | 80.01 | 58.38 | 69.25 |
|  | mean ROC-AUC | mean F1 | mean Bacc |
| AHLF | **85.98** | **74.25** | **76.06** |
| PhoStar | 80.99 | 61.32 | 72.95 |
|  | var ROC-AUC | var F1 | var Bacc |
| AHLF | **0.85** | **4.88** | **0.93** |
| PhoStar | 2.45 | 8.33 | 2.73 |

AHLF showed a better performance on average compared to PhoStar [11] in the detection of spectra of phosphorylated peptides (for evaluation details see Methods section). For example, AHLF achieved a higher median ROC-AUC than PhoStar (88.48 vs. 80.01) while also showing lower variances (AHLF 0.85 vs. PhoStar 2.45). Similary, AHLF outperforms PhoStar on the F1 and Bacc metrics. Note that the individual data sets vary greatly in size (number of spectra is ranging between a few spectra, e.g. 45 for ACHN up to 4,418,808 in case of the HeLa data set) and especially the combination of smaller data sets reflects diverse experimental setups. To check if performances depend on data set size, we calculated the overall metrics (by ignoring that spectra are organized in individual data sets) for the prediction scores from the hold-out folds (table 2). The overall metrics predominantly reflect the performances on larger data sets. Interestingly, both AHLF and PhoStar seem to perform better on larger data sets. This effect is less prominent for AHLF, e.g. the difference between its overall ROC-AUC and median ROC-AUC is only 3.54 but as high as 11.67 in case of PhoStar. This suggests that AHLF is more robust even with smaller data sets. Furthermore, some data sets seem to be more challenging to be characterized by either of the two approaches. We investigated if the performances of either tool are correlated across the different data sets. For example, the Spearman correlation coefficient between both approaches over the different data sets is 0.864 for the ROC-AUC score (supplementary material Fig. S2.1A). AHLF and PhoStar are different algorithms (neural network versus random forest-classifier), hence the correlation suggests that some data sets indeed tend to be more challenging than others.

**Table 2:** Overall balanced accuracy, F1-score and ROC-AUC as calculated over all hold-out predictions without averaging on the level of cell/tissue-types.

| Method | overall ROC-AUC | overall F1 | overall Bacc |
| --- | --- | --- | --- |
| AHLF | **92.09** | **85.51** | **83.68** |
| PhoStar | 91.68 | 79.89 | 82.00 |

Finally, we validate these findings by performing predictions on other phosphoproteomic data sets. We collected five recently published data sets containing non-human samples. AHLF was trained on data from human samples only, see above. Furthermore, the validation data resembles spectra that stem from four different instrument types as specified in the supplementary table S1.4.

AHLF performs better in 4 out of 5 data sets, specifically the ROC-AUC is 9.4 percent higher on average (table 3). PhoStar reaches its highest performance on a particular data set (PXD013868) with an ROC-AUC of 0.98 while AHLF achieves a comparable ROC-AUC of 0.95. AHLF seems to be more robust overall as its lowest ROC-AUC reads 0.79 but is down to 0.61 for PhoStar on JPST000703.

**Table 3:** Validation on recently published phosphoproteomic data. Bacc, F1-score and ROC-AUC for AHLF (A.) and PhoStar (P.).

| Data set | Bacc<br>(AHLF\|PhoStar) | F1-score<br>(A.\|P.) | ROC-AUC<br>(A.\|P.) |
| --- | --- | --- | --- |
| JPST000685 | 0.73\|0.57 | 0.71\|0.40 | 0.83\|0.66 |
| JPST000703 | 0.71\|0.55 | 0.67\|0.31 | 0.79\|0.61 |
| PXD013868 | 0.87\|0.93 | 0.94\|0.97 | 0.95\|0.98 |
| PXD014865 | 0.87\|0.81 | 0.88\|0.77 | 0.94\|0.90 |
| PXD015050 | 0.77\|0.66 | 0.77\|0.59 | 0.87\|0.76 |

### Interpreting AHLF reveals its ability to distinguish fragment- from noise-ions

Here, we investigate if AHLF is basing its decision on parts of the spectrum that belong to actual fragment-ions (peptide-related peaks) or rather on peaks that are considered noise-ions (ions that are not explained by the underlying peptide). Therefore we consult the original data base search results, which assigned a peptide to each identified spectrum. We compare those ground truth fragment-ions against peaks that appear important to AHLF for each spectrum individually. Hence, we calculated peak-level feature importances (SHAP-values) for individual spectra and compared them to the matching ions from the identified peptide (according to database search results). This is shown for a specific spectrum in Fig. 3A. For a quantitative comparison, we calculate a SHAP-value ratio, which is the sum of SHAP-values of matched ions divided by the sum of all SHAP-values. Intensity ratios were calculated accordingly. Both types of ratios are illustrated in Fig. 3A and explained in the Methods section. A visual inspection of the SHAP-value ratio versus intensity ratio for all spectra in the HEK293 data set indicates that SHAP-value ratios seem to be overall higher than their intensity-based counterparts (Fig. 3B). Meaning that AHLF can separate between fragment- and noise ions.

**Figure 3:**
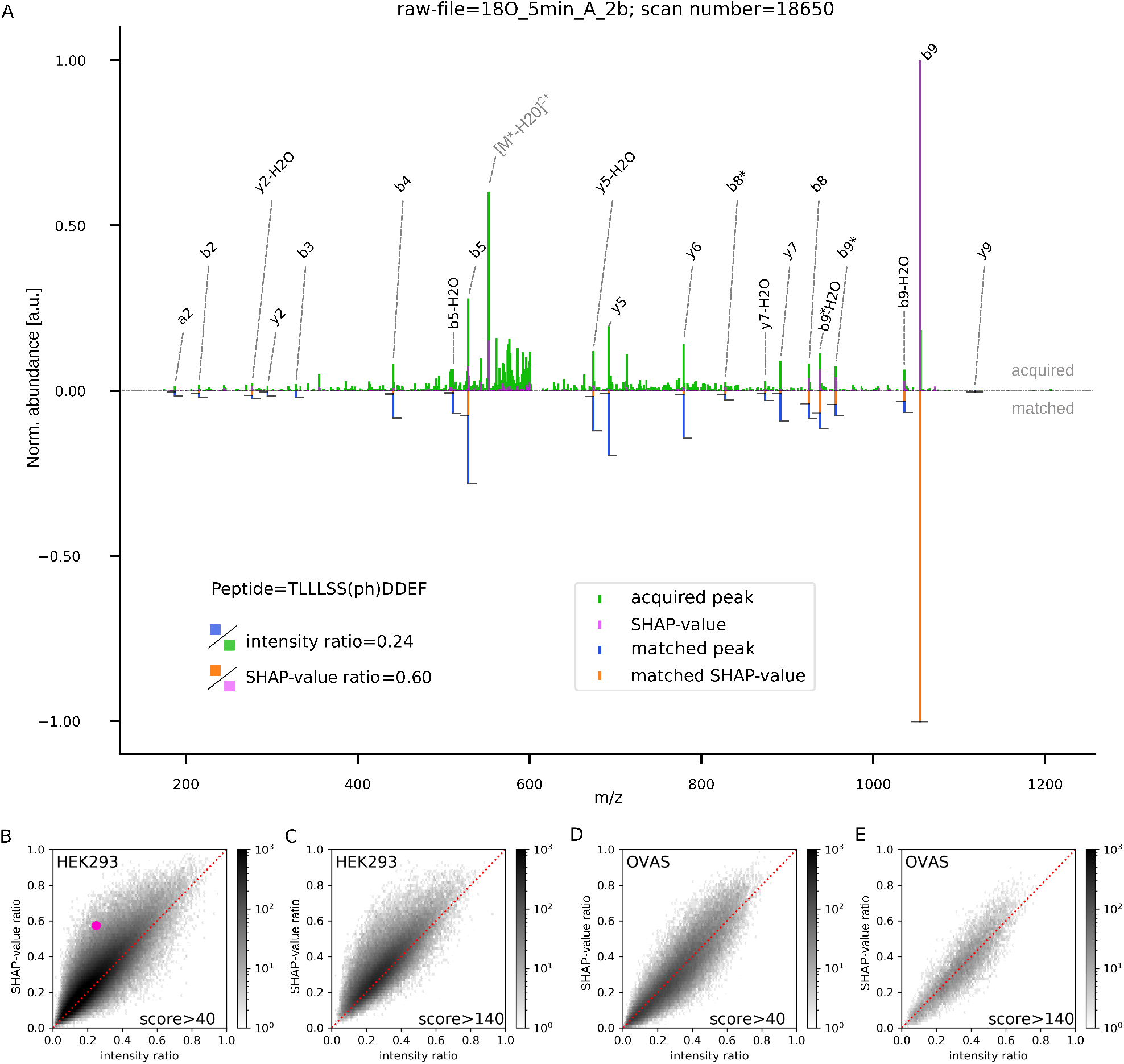
AHLF can distinguish between fragment- and noise-ions. **A:** mirror-spectrum drawn from the HEK293 data set as an example; positive values are acquired intensities (green) and SHAP-values from AHLF (magenta). Respectively, negative values are matched ions (blue; according to the database search results) and the matched SHAP-values (orange; subset of SHAP-value that are a the m/z-positions of matched ions). For a better comparison between matched intensities (blue) and each corresponding SHAP-value (orange) their heights are marked with whiskers. On the bottom left, the nominator and denominator for each of the two ratio types is indicated. Matched ions are labeled as annotated in the maxquant search results (we manually annotated the matching precursor ion [*M*^*^ – *H*_2_*O*]^2+^, which was not considered in the original search). **B-E:** hexbin plot comparing SHAP-value ratios to intensity-ratios as explained in the main text and illustrated in **A.** Each hexbin stands for a collection of ratios from a set of spectra, the number of spectra in each hexbin is given as logarithmic grey-code. The pink dot in **B** (SHAP-value ratio=0.60 and intensity ratio=0.24) indicates the bin which contains the example spectrum shown in panel **A. C:** same as **B** but spectra with lower scores(≤140) are removed. **D+E:** same as **B+C** but for the OVAS data set as a counter-example, where the SHAP-value ratios are not statistically higher than the intensity ratio (see Wilcoxon test below), which is not obvious from visual inspection alone.

This separation between signal and noise is not equally prominent throughout different data sets, for example in the OVAS data set it is less obvious and therefore shown as a counterexample in (Fig. 3D+E). In order to investigate if this signal-versus-noise separation appears throughout data sets, we performed a Wilcoxon signed-rank test on the 25 data sets from the first fold from the cross validation splits (fold a and the respective model AHLF*α*, see supplementary tables 1.2+1.3).

According to the Wilcoxon test for 18 out of 25 data sets the SHAP-value ratios are statistically higher than intensity ratios, considering a one-sided significance level of *α* = 0.01 (Bonferroni corrected). This supports our observation of AHLF learning a signal versus noise abstraction that is correct in most data sets.

Furthermore, we noticed that if the quality of the matching peptide is low (i.e. number and/or ion-current of matched ions is low) the SHAP-values falling onto these matching ions tend to be disproportionally smaller. This could mean that such peptides are false identifications from the database search. This is due to the fact that in MS-based proteomics a peptide is considered as identified, when the corresponding score of a that peptide-spectrum-match (PSMs) is above a selected level of an estimated false discovery rate (FDR), a typical FDR-level is 1% (Methods section). Hence, we perform comparisons of ratios for higher score thresholds (excluding spectra that have a lower Andromeda score [37]) (Fig. 3D+E). For six score thresholds ranging from 40 to 140 we performed Wilcoxon signed-rank tests (Fig. 4A). This illustrates the importance of PSM quality for model interpretation. Indeed, the higher score threshold diminishes the cases that pass testing for SHAP-value ratio being smaller than intensity ratio, (Fig. 4A, orange bars). In comparison, testing with the opposite alternative hypothesis is not impacted by higher score thresholds. Overall, this implies that AHLF was able to recognize the true fragment-ions in occasions, where a database search assigned potentially false identified peptides.

**Figure 4:**
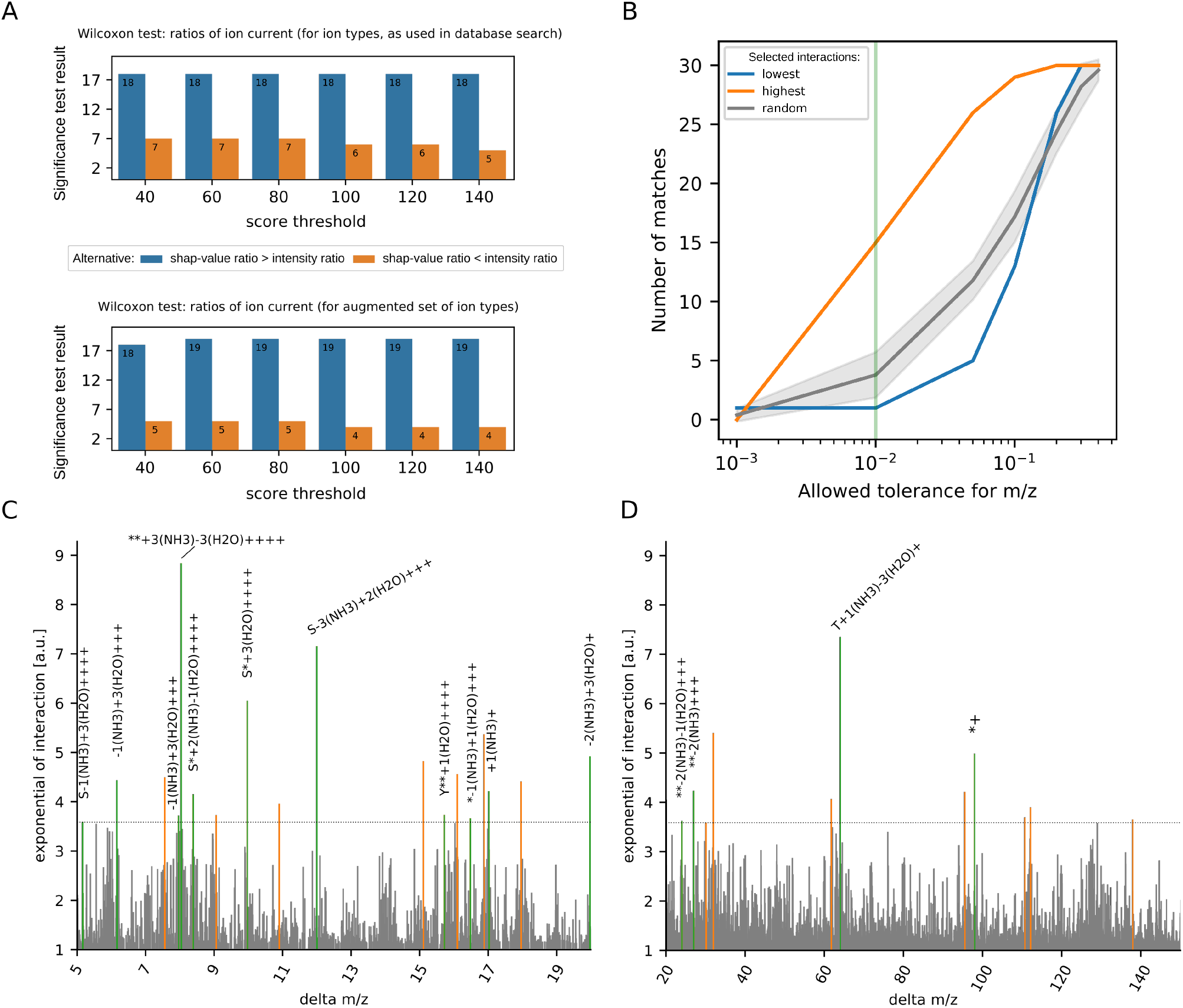
AHLF recognizes fragments beyond the classical b-/y-ions including biochemically relevant peak associations. **A:** number of data sets that pass the Wilcoxon signed-rank test depending on the applied score-threshold for the two alternatives: SHAP-value ratio>intensity ratio (blue) or SHAP-value ratio<intensity ratio (orange) for ion-types, as used in the database search (top) and using the augmented set of ion-types (bottom). **B:** adjusting the maximally allowed tolerance to 0.01 m/z (green vertical line). Comparatively, the top-30 highest interactions (orange) and potential negatives are estimated by i) the bottom-30 lowest interactions (blue) and ii) randomly sampled (grey; mean and standard deviation) of size 30 interactions are shown. **C+D:** pairwise interactions incorporate a combination of S/T/Y, phosphoric acid (*) and potential loss of ammonia (*NH*_3_) and/or water (*H*_2_*O*). Top-30 of highest interactions (above dashed line) of which 15 meet the required tolerance (green) are explained as annotated. The remaining 15 interactions (orange) do not meet the required tolerance even though still reflect reasonable interactions, see supplementary material. (**C+D** show the same data but were separated into two panels for a better readability and zoom into delta m/z between 5 and 20 m/z. Only the highest ten thousand interactions are shown here.)

Besides the quality of PSMs, the considered ion types determine which peaks are considered as fragment- or noise-ions. In the original database search, only nine types of ions for different charge-states are considered. Interestingly, in the example spectrum in Fig. 3A the most abundant peak [*M*^*^ – *H*_2_*O*]^2+^ is a doubly charged precursor ion subjected to a neutral loss of phosphoric acid and water (a precursor ion reflects an intact peptide). This particular combination of precursor ion and losses is originally not considered by the MaxQuant search engine but we rather annotated it here for illustration purposes. Obviously, AHLF considers this particular ion to be important, as can be seen from the SHAP-value assigned. This precursor ion even renders the second most important feature in this spectrum. Hence, we performed the feature analysis again but included additional ion types (Methods section).

When considering the augmented set of ion types, in up to 19 (out of 25) data sets the SHAP-value ratios are significantly higher than their intensity-derived counterparts (Fig. 4A). Increasing the score threshold diminishes the cases in which AHLF was unable to pick up the fragment-ions correctly from 5 down to 4 out of 25 data sets. Altogether, in the tested situations AHLF was able to separate fragment- from noise-ions for a vast pool of spectra in the majority of data sets and even does recognize ion-types, which where not considered in the original database search. This highlights the power of AHLF, resulting from the end-to-end learning paradigm, which leads to the detection fragmentation patterns reflecting existing structure within the spectra instead of biases introduced by prior knowledge.

### Pairwise interactions suggest that AHLF recognizes biochemically relevant associations between peaks

The ability of AHLF to distinguish between signal and noise is essential but it does not show whether AHLF has gained further mechanistic insights. Here, we investigate the recognition of biochemically reasonable patterns, but feature importance measures like SHAP-values are limited and do not reveal possible learned associations between features. Hence, we compute pairwise interactions [35] in combination with delta m/z between any peaks. These naturally match the notion of neutral losses, fragmentation of the modifying molecule itself and the fragmentation of individual amino acids. These processes are reflected by pairs of peaks with a distance matching the molecular weight of the lost group. Therefore we collected the identified phosphopeptide spectra from an individual run of the HEK293 data set. We kept spectra, which AHLF predicted as being phosphorylated, and calculated pairwise interactions and respective delta m/z for each spectrum (Methods section).

At first glance, pairwise interactions reveal a set of distinctively higher interaction strength at certain delta m/z (Fig.4D+E). For example, a prominent interaction is at 98 m/z, which reflects phosphoric acid, see annotated interaction ‘*’ (Fig.4E). To identify the other interactions, we searched delta m/z including combinations of i) S/T/Y, ii) phosphoric acid (abbreviated with ‘*’), ii) loss of ammonia (*NH*_3_) and/or loss of (*H*_2_*O*) subject to charges between +1 to +4 (for detailed stoichiometry see Methods section). Hence, a total of 2,352 hypothetical delta m/z where tested against the top-30 highest interactions. Surprisingly, all 30 interactions where positively identified when using the instrument-specific error tolerance (Fig. 4B). Hence, we adjusted the tolerance according to two baselines i) bottom-30 lowest interactions and ii) randomly sampled interactions as these should have few or no matches (Fig. 4B). Indeed, a much smaller tolerance needs to be selected to keep potentially false positives under control. We settled for a tolerance of 0.01 m/z, as this still yields 50 percent identified interactions amongst the top-30 highest values. these are annotated in 4D+E and reflect complex combinations of neutral losses (compared to what is usually considered throughout a database search). Amongst the remaining unannotated interactions we could identify additional 11 interactions when using a slightly higher tolerance of 0.05 m/z, see supplementary material S2.3. For example, the common loss of water is additionally matched when allowing this higher tolerance.

### Transfer learning allows AHLF to detect cross-linked peptides

We demonstrated that AHLF is able to detect a PTM without relying on engineered PTM-related features but rather AHLF ad hoc learned relevant features on its own. As a follow-up, we investigated whether AHLF is able to perform distinct tasks, such as detecting cross-linked peptides. This is particularly interesting as these spectra contain information from two peptides and the cross-linker molecule. Hence, the apparent fragmentation patterns are not only different from the PTM-related ones but even more so daunting to engineer. Excluding the requirement to engineer these complex features would demonstrate the impact and broad scope of our approach. Additionally, the lack of labeled spectra on this particular task is challenging, hence we used transfer learning and thus re-used trained weights from the AHLF models above (Methods section). Currently, there is no method available for a spectrum-based detection of cross-linked peptide. Hence, we use our framework to train fully connected networks as baselines including the optimization of their hyperparameters.

We compare our approach to three baseline models that are comprised of fully connected networks that either got presented with: i) the two vector representation (*m/z*′,*I*), ii) a vector of only intensities (_,*I*) or iii) a vector of only mz-remainder values (*m/z*′,_). For each baseline model we did a random search for hyperparameters. For AHLFx we took our previously trained models AHLF*α*,*β*,*γ* and -*δ* (from cross-validation on phosphoproteomic data) and continued training on spectra that were identified as cross-linked peptides (or linear peptides for the negative class respectively). AHLFx has the same hyperparameters as AHLF but we manually increased dropout rate and learning rate as AHLFx overfits otherwise. Hyperparameter selection including early stopping was done on the random split in Fig. 5A. In comparison, performance on a holdout set, where BS3 as a different cross-linker was used, is shown Fig. 5B. We emphasize that any of these models (baseline models or AH-FLx) are a result of our framework altogether. Yet, we found the best performance could be achieved in an out-of-the-box fashion in case of AHLFx. In contrast, the baselines required extensive hyperparameter search in order to achieve shown performances.

Finally, we validate AHLFx on PXD012723 [33] with BS3 as cross-linker. AHLFx shows an ROC-AUC of 0.88, whereas the baselines have comparable performances with an ROC-AUC of up to 0.87 (table 4). Even though the baselines show a higher optimal 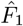 score of up to 0.71 we found that AHLFx better recognizes the positive class, which are spectra from cross-linked peptides. For AHLFx, the predictions scores of the positive class are well separated but mostly equally distributed in case of the fully connected networks. This is reflected by a better precision of 0.34 at a recall level of 0.95 for AHLFx. Whereas AHLFx indeed learned to detect cross-linked peptides the baseline models mostly only detect the linear peptides. Overall, this hints that AHLF has acquired a more general understanding of how peptides fragment, as transfer learning (carrying knowledge from phosphoproteomic to cross-link data analysis) helps AHLFx to detect spectra from cross-linked peptides.

**Table 4:** AHLFx detects spectra of cross-linked peptides as validated on PXD012723. ROC-AUC, F1-score and Precision at Recall of 0.95 are shown for AHLFx and fully connected networks as baselines.

| | ROC-AUC | $\hat{F}_1$ | Precision<br>(at Recall of 0.95) |
| --- | --- | --- | --- |
| AHLFx | **0.88** | 0.64 | **0.34** |
| Fully(m/z', l) | 0.87 | **0.71** | 0.20 |
| Fully(-, l) | 0.87 | 0.68 | 0.22 |
| Fully(m/z', -) | 0.72 | 0.58 | 0.19 |

## Discussion

We present a novel method for detecting PTMs and cross-linked peptides based on their fragmentation mass spectra without a database search. Our approach showed a high and robust performance over a wide range of data sets. We demonstrated that AHLF has learned substantial and fundamental fragmentation-related features, which was enabled by our strict end-to-end training schema. Interpretations of AHLF were in line with the majority of ground truth peptide identifications. Furthermore, AHLF in combination with SHAP was able to indicate potential false identifications, e.g. from database search results. This in-turn can highlight discrepancies in peptide-spectrum-matches (PSMs), which is at the core of almost any MS-based proteomics study. AHLF provides an unbiased and data-driven view on spectra and is orthogonal to currently existing MS-workflows, which often depend on prior knowledge such as data bases but also include heuristics on ion-types, losses and others. The combination of our results suggests that AHLF is expandable to perform other protein-related predictions based on peptide fragmentation spectra. We demonstrated this flexibility on cross-linking data, being the first time that this detection task was performed based on spectra alone and thus could speed up the computational analysis of cross-linking data.

For our approach we developed a specific two-vector representation of spectra. This allowed us to use convolutional layers and reduces memory-consumption without obscuring resolution-related information (Fig. 1). Overall, this enabled faster model training while making use of the original m/z values, as these can be recovered from mz-remainder, which is guaranteed by equation 1. Surprisingly, we could select a segment size that was around 18 times larger than the respective instrument specific resolution on that data set. The resulting two-vector spectrum accelerated training and subsequently we could train a more sophisticated model. Even though we adjusted the segment size based on specific data, the selected size not necessarily complies with all data sets. A smaller segment size might better suit experiments with spectra, where peaks are tightly packed, but this does not guarantee a better performance overall. Effectively, the segment size and window size are hyperparameters and affect the model capacity of AHLF. Similarly, others found that accounting for the resolution of mass spectra in the context of deep learning, in particular for *de novo* sequencing, is essential [38]. Finally, we empirically demonstrated the impact of our two-vector spectrum at a later stage in this work (Fig. 5).

In order to model fragmentation patterns we set up a deep neural network that we designed to promote learning of long-range associations between features (Fig. 2). Most biochemically related peak associations are long-range relations because peaks in a spectrum can be hundred Dalton (Da; atomic mass for a singly charged ion) or even multiple hundred Da apart (e.g. several phosphosites at different locations of a peptide). Similarly, in our two-vector spectrum representation related features can be multiple hundreds steps apart (Fig. 1–3). It is empirically known that recurrent neural networks (including LSTMs and GRUs) are unable to learn in such extreme regimes of long-range associations [39]. Hence AHLF falls into the category of temporal convolutional neural networks (TCNs), which are deep neural networks consisting of dilated convolutions and capable to learn long-range associations [39, 40].

With this approach, AHLF outperforms the current state-of-the-art PhoStar [11] on phosphopeptide detection on recently published data sets from diverse lab environments and experimental setups. In particular, we could demonstrate that AHLF is more robust against variables such as sample species, type of instrument and lastly data set size. The latter is an indirect measure of lab diversity which is mostly covered by smaller data sets where our model was showing a robust performance altogether (tables 1 and 2). Ultimately, our validation on unseen and recently published phosphoproteomic data confirms our results from the cross-validation benchmark and highlights the ability of AHLF to cope with the diversity between data sets. This suggests that AHLF is not a Clever Hans and our model did not just learn some shortcuts but rather recognizes phospho-specific pattern that are omnipresent throughout the various data sets.

Interpretation of AHLF and subsequently investigation of the mechanisms behind what AHLF has learned coincided with biochemically reasonable fragmentation patterns. In contrast to previous approaches, AHLF is a generic deep learning model and the abstraction of important features from spectra is fully integrated into the learning process itself. We investigated, after AHLF was trained, if AHLF is picking up reasonable features from a given spectrum. Therefore, we derived feature importance values on the level of individual spectra for a large collection of spectra. We could show that AHLF was using the entire spectrum (instead of picking up single peaks) to derive its predictions score, as prominent SHAP-values are often scattered over the entire m/z range (Fig. 3A). Furthermore, important features coincide with peptide fragment-ions. This was true for most spectra from the tested data sets (Fig. 4A). We could quantify that AHLF was actually learning a suitable abstraction of a given spectrum and thus being able to focus on fragment-ions despite it did not know the peptide sequence in advance.

We noticed that our interpretation efforts are partially hindered by potential errors in the ground truth peptide-spectrum-matches (PSMs) assigned from the original database search. Imagine a falsely-identified PSM being used for our assessment, we would conclude that AHLF is looking at noise. When in fact, AHLF may focus on the true fragment-ions (those that were not assigned due to a wrong PSM resulting from the database search). We could quantitatively show that a ground truth including PSMs of higher quality (higher scores) lead to a better agreement between SHAP-values and matching ions (Fig. 3A+B) and 4A. This could mean that these spectra contain more information or are less noisy, hence could be better identified from the search engine and predicted by AHLF. But it could also be a hint that AHLF is able to reveal potential disagreement between a spectrum and a potential false positive identification. Additionally, when comparing the two ion-type models in Fig. 4 (top versus bottom panel) the increase in tests passed for the SHAP-value ratios greater than intensity ratios suggests that AHLF is recognizing additional ions (from the augmented ion set), which were originally not considered through database search. We speculate that AHLF (in combination with feature importance like SHAP) could be used to perform a post-search correction of PSMs. So-called rescoring methods are a highly effective and commonly used approaches to improve identification yields from a search engine [16]. But instead of only including features of PSMs one could reconcile AHLF (by deriving SHAP-values) to gain predominantly spectrum-related information. Note that the opinion of AHLF is not predetermined by the selected database. AHLF purposefully learns an abstraction of spectra and can distinguish between fragment-ions and noise peaks, despite we never presented such information on the level of individual peaks to our model.

Following up on this, we computed pairwise interactions between any pair of peaks in a spectrum. This way we could show that AHLF has a fundamental understanding of how phosphopeptides fragment, coinciding with commonly known neutral losses such as phosphoric acid and combinations thereof (Fig. 4C+D). This is analog to engineered features which are used in Colander [10] and Phostar [11]. But in contrast, AHLF is ad hoc learning them from the data and here we checked, after training, if those coincide with common expert knowledge. For example, the fragmentation of a phosphopeptide often involves the loss of phosphoric acid *H*_3_*PO*_4_, which results in a delta mass of around 98 Da between pairs of ions with a single positive charge [41, 13] and indicated by ‘*’ in Fig.4D. Additionally, we could explain 15 out of the top-30 highest interactions when matching delta masses with an allowed tolerance of only 0.01 m/z (Fig. 4B) and up to 26 when choosing a slightly higher tolerance of 0.05 m/z (Fig. S2.3). Conversely, we missed out on phosphoric acid undergoing an additional loss of water (a common phospho-specific loss at around 80 m/z). Despite, there is a substantial interaction recognizable at 80 m/z it is not amongst the selected top-30. Furthermore, we observe reasonable but rather complex combinations of losses for multiple charged ions. As these should be less common, we double checked on that by performing the same kind of analysis but for data, which had been acquired on an Orbitrap instrument involving a higher resolution mass analyzer (Fig. S2.4). This allowed us to de-isotope and de-charge these spectra, which summarizes all isotopic and possibly multiple charged peaks into a common mono-isotopic peak. On the one hand side, multiple interactions were accumulated into fewer peaks such that the dynamic range between potentially false and true interactions was reduced and we had to select the top-500 highest interactions to match reasonable interactions (such as 98 Da for phosphoric acid for example). On the other hand side, our hypotheses were reduced by factor 4 down to 588 because there was no need to account for the different charge states. This reduces the potential to match false positive delta masses. Additionally, we were able to select a much smaller allowed tolerance of only 0.001 Da under which we could match 48 interactions values (for 16 unique delta masses). As a result, we again observed delta masses that involved combinations of multiple losses of water and/or ammonia. Overall, delta masses (respectively delta m/z for iontrap data above) for high interactions values match our hypotheses, whereas randomly selected interaction values or those for low interaction values do not match our hypotheses. Keep in mind that the masses are existing in the spectra anyways and matching the delta masses alone does not tell us much about AHLF. But AHLF accounts for the interaction values and the higher ones were more likely to be explained (Fig. 4A, S2.3A and inlay in Fig. S2.4). In conclusion, we observed rather complex combinations of neutral losses, which included multiple losses of either water and/or ammonia. This is partially in contrast to what was considered in PhoStar [11]. Even though also both types of losses are considered in their random forest model they miss out on combinations involving multiple losses (up to three) of either type and mixtures thereof. Overall, AHLF has indeed ad hoc learned task-specific fragmentation even though it has never explicitly received information on that level, e.g. annotated peaks were not provided as an input to AHLF.

Note that we did not use any further knowledge from the database search results to identify these losses (except the information that we look at identified phosphopeptides only). This implies that, in principle, one could identify and propose new fragmentation patterns even for other post-translational modifications or even other less related tasks.

Even though we limited ourselves to the interpretation of feature importance and pairwise interactions this does not mean that AHLF is not learning more complex patterns. A spectrum is comprised of internal fragments [42], complex isotopic distributions and further fragmentation of post-translational modifications themselves [12]. This results in a much more complex picture when all possible associations are considered. But there is no conceptual hindrance for AHLF to learn these complex patterns and our model has the following advantages: (i) it is not limited to a fixed set of ion types, (ii) it does not exclude internal fragments and neutral losses, (iii) it is possible to learn non-proteinogenic amino acids [18] and to adjust the alphabet size accordingly [43], (iv) it allows unrestricted study of post-translational modifications and (v) it is orthogonal to a database search and thus can reduce the search space or reveal discrepancies in the search results.

In fact, we demonstrated a broader applicability of our framework by using transfer learning. This is crucial for situations where training data is limited, e.g. for cross-linked peptides we had thousands of annotated spectra whereas we gathered millions of annotated spectra for the phosphoproteomics approach. A cross-linked peptide detection potentially reduces computational costs in a dedicated cross-link database search, where the search space grows quadratically with candidate peptides, as any pair of peptides needs to be searched [14]. For the detection of cross-linked peptides we could show that AHLFx is able to perform better than our baselines. For the latter we had to perform expensive hyperparameter estimation. In contrast, AHLF became AHLFx in an out-of-the-box fashion by simply continuing training on the new task with pre-trained weights and already existing set of hyperparameters. Based on this, we anticipate that besides transfer learning an even simpler finetuning scheme, e.g. to a specific lab environments using in-house data is feasible. This would help to overcome the data inherent variability which we observed in table S1.5 and potentially boost the performance of AHLF in lab-specific workflows.

When proposing the two-vector representation we primarily focused on its technical aspects. Yet, the mz-remainder as a feature is related to the so-called fractional mass (a special case of mz-remainder, when segment-size is 1 m/z) depends on the elemental composition of a peptide [44]. Hence, mz-remainder is valuable information about fundamental properties of the peptide and can thus support the detection of a modified peptide. We investigated the impact of keeping the mz-remainder (Fig.5) and found that including it as a feature substantially improved the detection of spectra from cross-linked peptides.

**Figure 5:**
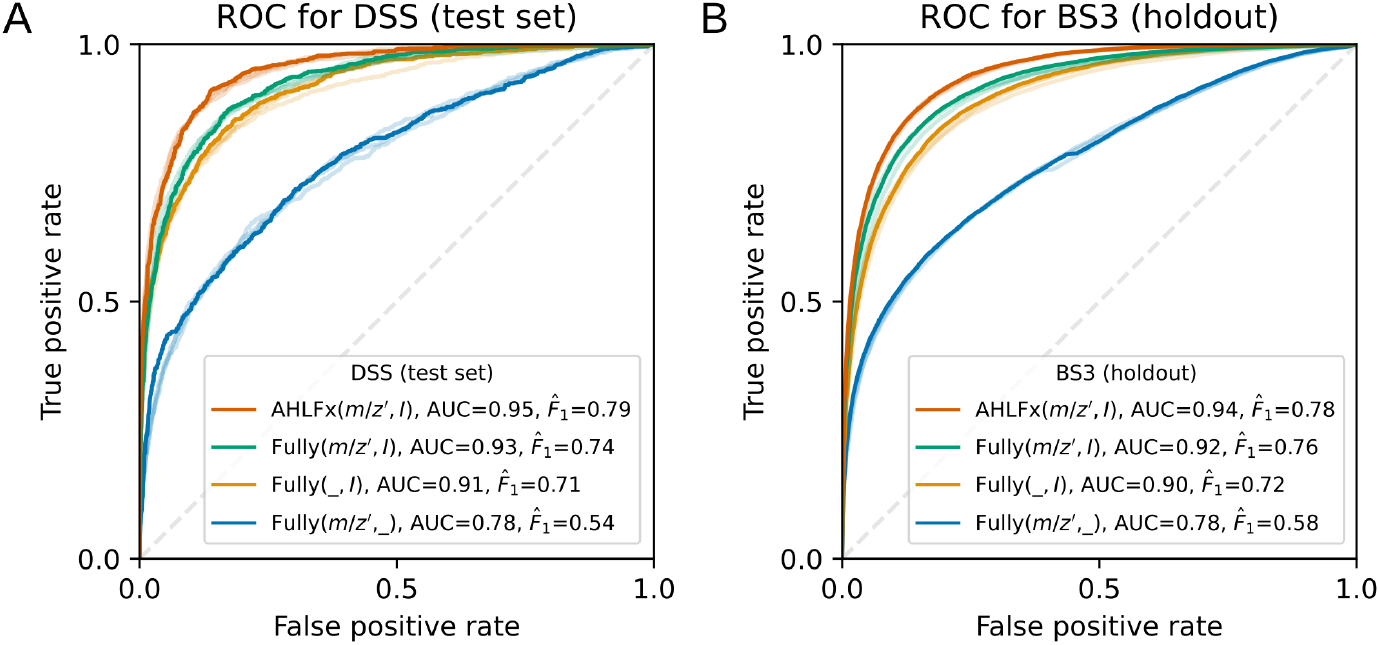
AHLF detects cross-linked peptides via transfer learning. **A:** evaluation on the test set for the DSS cross-linker and **B:** holdout set for the BS3 cross-linker. Receiver operating characteristic of AHLFx (orange) compared to three fully connected networks as baselines (green, yellow and blue) and a random baseline with ROC-AUC=0.5 (dashed line). For each, the top-4 training outcomes are shown and the best is highlighted respectively. For each best model, ROC-AUC and optimal F1-score are summarized in the legends.

In analogy, we observed that the number of potential false identifications of delta m/z depends on the allowed mass tolerance (Fig. 4B). This suggests that AHLF makes use of the available information that comes from keeping the mz-remainder and withholding the mz-remainder from AHLF would probably negatively impact its performance.

We foresee that our approach will spark diverse future applications since feature creation is fully integrated in the learning process. Possible future applications include the prediction of phosphosite localization scores, spectrum-identifyability scores, post-search re-scoring, biomarker detection and anomaly detection including the detection of non-proteinogenic amino acids and uncommon PTMs. To the best of our knowledge, this is the first deep learning model being able to ad hoc learn fragmentation patterns in high-resolution spectra. Additionally, our specific set of hyperparameters may serve as a starting point and speed up hypothesis testing.

## Methods

### Public data sets used for cross-validation

For training and testing through cross-validation, we used the combined data set PXD012174 [4], containing 112 individual repositories organized according to 101 human cell/tissue types (number of zip-files in PXD012174 that contain fragmentation spectrum raw-files). The individual data sets used phospho-enrichment assays. This combined data set was re-analyzed by Ochoa and colleagues and did undergo a joint database search using MaxQuant [45] with the following error rates: false-discovery rate (FDR) was set to 0.01 at peptide-spectrum match (PSM), protein, and site decoy fraction (PTM site FDR) levels. The minimum score for modified peptides was 40, and the minimum delta score for modified peptides was 6. The combined search results were taken from the ‘txt-100PTM’-search results, as described in PXD012174 [4]. For each spectrum, a label (phosphorylated or unphosphorylated) was assigned when the set of PSMs exclusively contained phosphorylated or un-phosphorylated peptides. In other words, if the set of identifications for a given spectrum contained PSMs for both kinds of peptides (phosphorylated and unphosphorylated), then we discarded the spectrum. This way, we try to avoid ambiguous labels. Overall, this yields a training set that contains 19.2 million PSMs consisting of 10.5 million phosphorylated PSMs and 8.7 million unphosphorylated PSMs.

### Model training

We optimized the cross entropy loss using the adaptive learning rate optimizer ADAM [46] with an initial learning rate of 0.5 · 10^−6^. AHLF was trained for 100 virtual epochs consisting of 9000 steps, which was a gradient decent step on a mini-batch of size 64. For regularization we used early stopping and drop out with a rate of 0.2 on the fully connected layers. We initialized weights for all layers using ReLU-activation with random weights drawn from a standard normal distribution using He correction and Glorot correction for layers, not using ReLU-activation. See supplementary material for details on hyperparameter optimization.

### Public data sets used for validation

For validation we use five data sets [28, 29, 30, 26, 27]. For each data set, we used the original search results from MaxQuant alongside the raw spectra. Additionally, we removed PSMs with scores smaller than 40. Detailed information are summarized in the supplementary table S1.4 for each data set.

### Evaluation of AHLF

We compared AHLF against PhoStar [11], currently state-of-the-art method for phosphopetide prediction based on fragmentation spectra. We used PhoStar with default parameters: m/z tolerance was set to 10 ppm, peak-picking depth was set to 10 (per 100 m/z) and score threshold set to 0.5 for all results shown here. PhoStar is closed source and not trainable by the user. We used the original PhoStar ensemble model parameters [11]. For 462,464 spectra from PXD012174 (2.4% of the data set) PhoStar was technically not able to predict a classification score (it provides an error about mis-matching masses). This also applies to some spectra from the validation data. We proceeded by assigning a PhoStar-prediction score of 0.5 to these spectra in order to achieve a fair comparison anyways.

To calculate metrics such as balanced accuracy, F1-Score and the area under the (receiver operating characteristic)-curve (ROC-AUC) we used Scikit-learn [47]. We set a score-threshold to 0.5 to get class labels from the predicted continuous binary-classification scores.

In case of four-fold cross-validation the four resulting models of AHLF were evaluated on their respective hold-out data set (supplementary table S1.3). For the benchmark on unseen, recently published data we used the arithmetic mean of prediction scores from the AHLF model ensemble.

### Methods used for explaining AHLF

For the calculation of feature importances we used SHAP [34]. From the SHAP framework, we use Deep-Explainer and we set an all zeros vector as background reference spectrum. In particular, we chose SHAP because it allows us to investigate each spectrum individually (in contrast to global methods which only report aggregated statistics over multiple data points). Furthermore, SHAP computes importance values that are additive, which means for a given spectrum their sum is supposed to mirror the prediction score of AHLF for that spectrum. In our case, the error between the sum of SHAP-values and AHLF scores were smaller than 1%. Hereinafter, we refer to the additive feature importance values as SHAP-values.

Furthermore, we are interested in absolute SHAP-values as we investigate both types of spectra (either from a phosphorylated or un-phosphorylated peptide) equally. In particular, we are testing whether AHLF can separate fragment- from noise-peaks. Therefore, we assume that for each spectrum the sum of all intensities Σ*I*≔ *I*_*matching*_ + Σ*I*_*non–matching*_ is consisting of intensities that are at an m/z that could be matched to a peptide fragment ion and others that could not be matched. Furthermore, we define an intensity ratio ≔Σ*I*_*matching*_/ Σ*I* and analogous SHAP-value ratio ≔Σ|s|_*matching*_/ Σ|s| Note, these ratios are bound between zero and one and any scaling factor is canceling out such that we can compare the two types of ratios easily. We anticipate that our explainabilty assessment depends on which ions are matching, hence we will investigate crucial parameters that potentially alter the ground truth via the resulting PSMs and used ion-types systematically below.

In order to have a ground truth on the level of each individual peak within a given fragmentation spectrum we used the PSM information from the search results. For our tests using the original set of ions (as matched during database search), we directly used the MaxQuant output from the msms-table. For each PSM, a set of matching peaks including m/z, intensity and ion-type has been reported. The ion types that were used during database search contained: a, b, y, b-H3PO4, y-H3PO4, b-NH3, y-NH3, b-H2O and y-H2O ions. In contrast, for our experiments with an augmented set of ion types, we additionally computed the theoretical spectrum for each peptide for a given PSM. Therefore, we augmented the above listed ion types by including these types: a-H2O, a-NH3, c, cdot, c-H2O, c-NH3, M, M-H2O, M-NH3, x, x-H2O, x-NH3, z, z-dot and z-H2O. We made sure to reproduce the fragment masses that were matched during the original search and concatenated the additional calculated fragment-ions yielding an augmented theoretical spectrum. In order to find matching peaks between an acquired and the augmented theoretical spectrum, we used a binary search as implemented by [48]. For matching peaks we allowed a mass tolerance of either 0.5 Da, in case of an ion trap, or 20 ppm, in case of an Orbitrap.

For each spectrum we are able to compute two types of ratios as stated above (both reflecting a measure of signal-versus-noise). We assume that these resemble two random variables that are measured on the same spectrum. Therefore, we chose the Wilcoxon signed-rank test to compare whether one ratio is statistically greater than the other ratio. We chose a significance level of *α* = 0.01 for the one-sided signed-rank test and used Bonferroni correction to account for the number of thresholds times number of datasets as total number of hypotheses. Test statistic and p-values were computed using the Wilcoxon test from Scikit-learn [47].

For pairwise interactions we used Path Explain [35]. Path Explain computes interactions between any pair of input features for a deep learning model, in our case any pair of peaks within a spectrum. Computationally, this is very expensive (more than 1 hour per spectrum). We excluded spectra with more than 500 peaks (Path Explain computations increase quadratically with number of peaks). We selected a single run (‘18O_5min_A_2b’ from the HEK293 data set). Furthermore, we selected spectra from identified phosphorylated peptides (by MaxQuant) and required AHLF to correctly predict them, any other spectrum was discarded. This yields 193 spectra for which we computed pairwise interactions. In case of the Orbitrap run, we used (‘5_min_M_a_QE.raw’ from the HeLa data set). Again we kept spectra for identified phosphorylated peptides only, resulting in 504 spectra for which pairwise interactions are calculated. In any case, we kept positive interactions as we are interested in the positive class and the responsible pair-wise interactions. In case of the Orbitrap run, we used the python package ms-deisotope to de-charge and de-isotope the spectra. For each isotopic envelope (groups of peaks are summarized as one monoisotopic peak) we averaged over the pairwise interactions by using the arithmetic mean.

To identify the interactions and assigned neutral losses, we searched delta m/z by including any combination of i) [0,1] of S/T/Y, ii) [0,+1,+2] of phosphoric acid (abbr. with *), ii) [−2,−1,0,+1,+2] losses of ammonia (*NH*_3_) and/or [−2,−1,0,+1,−2] losses of (*H*_2_*O*) subject to charges between +1 to +4. These combinations resemble 2352 different delta m/z. In case of the Orbitrap data, the number of hypotheses was reduced to 588, because there was no need to account for different charge states. In the main text we refer to a peak-matching-equivalent tolerance of 0.01 m/z.Since delta masses are matched, we multiply this tolerance by 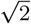 i.e. according to Gaussian error propagation for a difference of *mz*_1_ − *mz*_2_, the effective error is 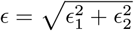 such that we used 0.01· 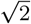 as apparent tolerance when matching delta m/z.

### Transfer learning on public cross-link data

For training and testing we utilized public data from JPST000916 (DSS as cross-linker) and for evaluation we used JPST000845 (BS3 as cross-linker) as hold-out set [31, 32]. Raw files were converted to MGF and after m/z recalibration searched with xiSEARCH[49]. As labels we used cross-linked peptide spectrum matches (CSMs) from xiSEARCH results as positive class (at 5% CSM FDR). And the reported linear PSMs as negative class respectively (5% PSM FDR). During transfer learning we took our pretrained models AHLF*α*-*γ* and continued training on JPST000916, we adjusted learning rate to be 0.0001 and increased the dropout-rate accompanying the dense-layers to 0.5. All other hyperparameters were kept the same as for the original training of AHLF. For the baseline models we used a fully-connected network with number of layers=2,3,4,5 or 6 with number of units = 32, 288, 544, 800 or 1024 with a dropout-rate of 0.0, 0.2 or 0.5 using ADAM with initial learning rate from 1.0·10^−3^, 0.5·10^−3^,1.·10^−4^,0.5·10^−4^ to 1.0·10^−5^. We randomly sampled 50 parameter-constellations and trained three times each. From the resulting 150 training runs we evaluate the best 4 runs. This was repeated for the feature vector (*m/z*′, *I*), and the two masked-out versions (_,*I*) and (*m/z*′,_), where respective values where set to zeros. During transfer learning and baseline-training we used early stopping on the test set split.

### Spectrum representation of AHLF

With the help of our particular spectrum representation we exploit the sparsity of centroided spectra. A centroided fragmentation spectrum is a list of peaks that are tuples of mass-over-charge (*m/z*) and intensity (*I*). If the sparsity matches the segment size (meaning exactly one peak per segment) this operation is reversible as illustrated in Fig. 1A. In other words, one is able to recover the original peaks-list from our two-vector representation. To achieve a truly loss-less conversion we can exploit the sparsity of centroided spectra and adjust the segment size accordingly. Theoretically, the chance of two peaks randomly falling into the same segment is marginal. A peaks-list contains *l* total number of peaks. Typically, a ms/ms spectrum has hundreds, in extreme cases up to a few thousands of centroided peaks *l* and a dense vector matching the instrument resolution has easily multiple hundreds-of-thousands of entries *b*. Assuming uniformly sampling of values for m/z, the probability for randomly choosing a m/z (within the resolution) that has been occupied already is given by *1*/(*b − l*). It is known that m/z displays dead spots (e.g. combined histogram of m/z from all peaks-lists). Hence, m/z is usually not uniformly distributed. However, the opposite case of a fragmentation spectrum where all peaks are clumped together is usually discarded during quality assessment. These poorly fragmented spectra are not very informative.

In the Results section, we describe a strategy of how to generate our proposed spectrum representation from a given peaks-list (Fig. 1A). An alternative, but equivalent strategy is to (i) populate an all-zero vector (of size window-range times instrument-resolution) with peaks and (ii) apply the maximum and arg-maximum within small mass segments of fixed size (Fig. 1A). Furthermore, our spectrum representation reflects a regular grid, which is equally spaced and with fixed connectivity of m/z-values. This makes a spectrum compatible with a network that uses convolutional or recurrent layers (comparable to applications including image or a time-series with constant time-steps).

To handle the amount of spectra, especially for feeding a GPU with training samples, we set up a custom pipeline facilitating the conversion of peaks-lists into the two-vector representation. It also performs common preprocessing steps, e.g. ion-current normalization that is dividing intensities by the sum of squared intensities. Our preprocessing pipeline is implemented in Python, largely integrating Pyteomics [50, 51] and tensorflow [52]. The combined preprocessing and training pipeline can be found as part of our code repository. Our pipeline accepts mgf-files as input-files. Spectra from raw-files were centroided and converted to mgf-files by using ThermoRawFileParser [53].

### Details about particular choices for the model architecture of AHLF

In the Results section, we illustrate the model architecure of AHLF (Fig. 2). We show how a single output (black arrow, center) can receive information from any input (black arrows) via a collection of paths (black solid lines) in the block of dilated convolutions. The block of dilated conolutions facilitates a receptive field that spans the entire feature vector. The block has nested stacks of convolutional layers and 64 filters per layer, hence the actual model complexity is not fully captured by the simplified illustration here (blue inlay, Fig. 2). On the right in Fig. 2 we compare what is drawn versus what was implemented (in parentheses). Additionally, we illustrate how a single output can learn to reflect a specific pair of ions (pink path in Fig. 2). Associations between more than two input features can be learned by the model, but not illustrated here. Furthermore, we introduced skip-connections by using convolutions with kernel-size=1 that are added to the stacked convolutions (blue inlay on the right in Fig. 2). This allows for inputs to be passed to the output (yellow path in Fig. 2).

A block of dilated convolutions is commonly called temporal convolutional neural network (TCN) [39]. In a TCN, the receptive field grows exponentially and therefore the gradient computation only needs log(distance-between-features) steps. We preferred this over, e.g. a transformer architecture [54], even though the latter facilitates gradient computation that are independent of the distance between two features. But it requires to keep a self-attention matrix, which in-turn scales quadratic with input-size. In contrast, for a TCN, the computational complexity of convolutions scales linearly with the input-size [54]. Hence, in case of AHLF we chose a TCN over a transformer.

In our TCN, we use padding that conserves the size between the input and the output layer (so-called ‘same’-padding). In contrast, TCNs are also usually used in conjunction with ‘causal’-padding, e.g. the ‘Wavenet’ is using causal-padding [55]. This is due to the fact that the latter is modeling audio signals over time, and feedback from the future to the past is eliminated by ‘causal’-padding on purpose. In case of a fragmentation spectrum, the peptide fragmentation happens from both ends of a peptide and a spectrum is bi-directional in that sense, hence we decided to stick with ‘same’-padding. An output feature is able to receive information from the left and the right part of the previous layer (Fig. 2). The TCN-block is followed by fully-connected layers. The final prediction is facilitated by a sigmoid as activation-function, which outputs a score between zero (unphosphorylated) and one (phosphorylated).

## Code availability

An open source implementation with command-line instructions is publicly available (under MIT licence) at https://gitlab.com/dacs-hpi/ahlf and includes four independently trained models. Additionally, the code-repository allows fine-tuning of AHLF as well as training from scratch on third-party or user-specific data.

## Acknowledgements

This work was supported by the Deutsche Forschungsgemeinschaft (grant RE3474/2-2 to BYR) and by the BMBF-funded de.NBI Cloud within the German Network for Bioinformatics Infrastructure (de.NBI) (031A537B, 031A533A,031A538A, 031A533B, 031A535A, 031A537C, 031A534A,031A532B). We thank Philipp Benner, Tobias Loka and Elizabeth Y. Yuu, for proofreading this manuscript.

## Supplementary material

**Algorithm 1.**
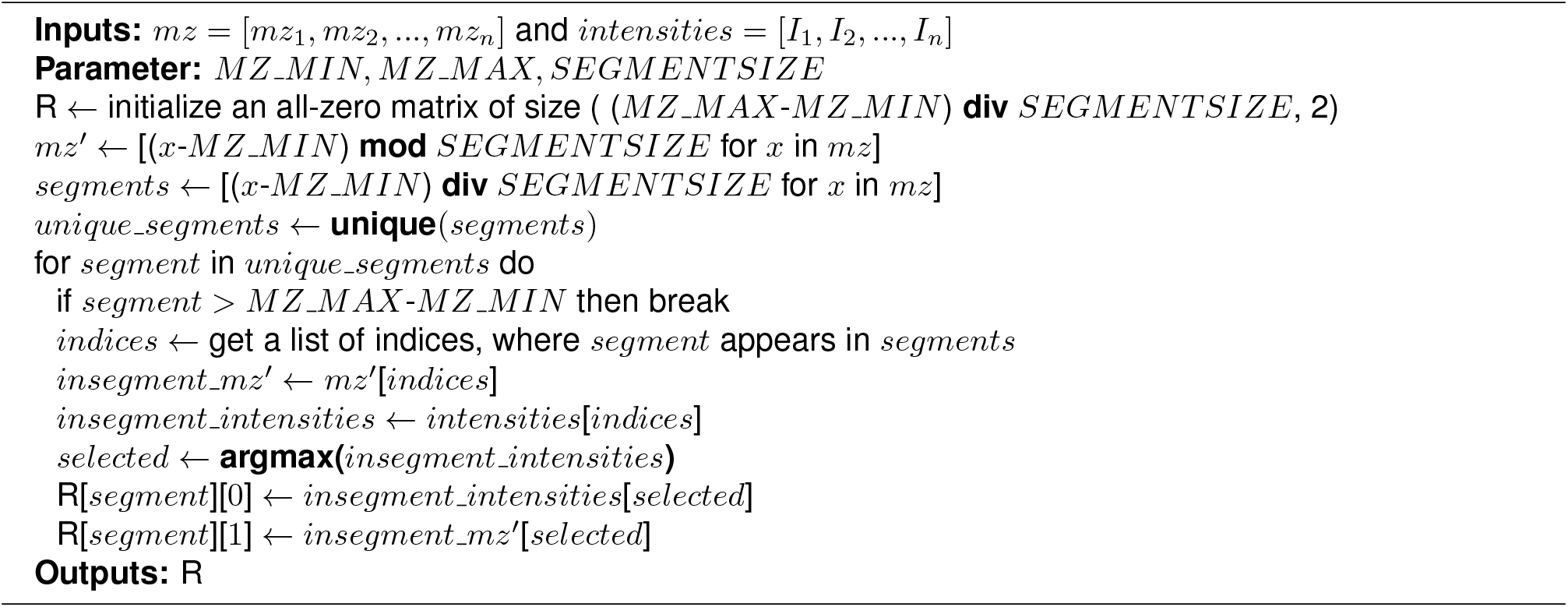
converts a peaks-list into a two-vector spectrum representation containing mz-remainder and intensities.

## S1 Supplementary tables

**Table S1.1:** Model architecture with 6,705,985 trainable parameters.

| type | n - filter (or units) | size | dilation rate | activation |
| --- | --- | --- | --- | --- |
| conv1d | 128 | 1 | 1 | ReLU |
| 2xconv1d/conv1d | 64 | 2/1 | 1 | ReLU |
| 2xconv1d/conv1d | 64 | 2/1 | 2 | ReLU |
| 2xconv1d/conv1d | 64 | 2/1 | 4 | ReLU |
| 2xconv1d/conv1d | 64 | 2/1 | 8 | ReLU |
| 2xconv1d/conv1d | 64 | 2/1 | 16 | ReLU |
| 2xconv1d/conv1d | 64 | 2/1 | 32 | ReLU |
| 2xconv1d/conv1d | 64 | 2/1 | 64 | ReLU |
| 2xconv1d/conv1d | 64 | 2/1 | 128 | ReLU |
| 2xconv1d/conv1d | 64 | 2/1 | 256 | ReLU |
| 2xconv1d/conv1d | 64 | 2/1 | 512 | ReLU |
| 2xconv1d/conv1d | 64 | 2/1 | 1024 | ReLU |
| 2xconv1d/conv1d | 64 | 2/1 | 2048 | ReLU |
| 2xconv1d/conv1d | 64 | 2/1 | 4096 | ReLU |
| maxpooling | - | 20 | - | - |
| dropout(rate=0.2) | - | - | - | - |
| fully connected | 512 (units) |  |  | ReLU |
| dropout(rate=0.2) | - | - | - | - |
| fully connected | 512 (units) |  |  | ReLU |
| dropout(rate=0.2) | - | - | - | - |
| fully connected | 512 (units) |  |  | ReLU |
| fully connected | 1 |  |  | Sigmoid |

**Table S1.2:** Four-fold cross-validation splits. Individual data sets in PXD012174, sorted by size (number of spectra; largest at the top).

| a | b | c | d |
| --- | --- | --- | --- |
| HeLa | Primary-Ovarian | Jurkat | HCT116 |
| HEK293 | A431 | DG75 | M019i |
| Primary-AML | BoneMarrow | SW480 | MCF7 |
| Daudi | A673 | Lung | Kit255 |
| U2OS | Mono-Mac-1 | MV-4-11 | SKM-1 |
| Primary-Gastro | WM239A | 143B.TK | BT474 |
| A549 | Fibroblast | U937 | Primary-Pancreas |
| HaCaT | HUVEC | H1975 | Metastasis-Pancreas |
| OVAS | KOC-7C | HCCLM6 | OVISE |
| Primary-Prostate | MCF10A | Primary-Melanoma | SH-SY5Y |
| TOV-21-Primary | Primary-BOEC | Primary-Colorectal | Platelets |
| LNCaP | MDA-MB-231 | H3255 | 11-18 |
| HEPG2 | Primary-Breast-MicroAndExo | H1299 | SK-N-BE |
| Colon | Primary-Glioblastoma | RKO | Brain |
| Kasumi-1 | Muscle | COLO-205 | P31.Fuj |
| HT-29 | SW1398 | THP1 | ECV-304 |
| ES2-Primary | DLD-1 | CACO-2 | GB2 |
| OVSAYO | H358 | Primary-Glioma | Primary-Liver |
| HDMVEC | DoHH2 | SU-DHL-6 | RL |
| PANC-05-04 | CTS | PC9 | Capan-1 |
| SU.86.86 | HPAF-II | BxPC-3 | PANC-04-03 |
| CFPAC-1 | PANC-08-13 | Capan-2 | SW1990 |
| RPMI-8226 | Hs-700-T | PANC-10-05 | PL45 |
| HPAC | Hs-766-T | PANC-03-27 | OMP2 |
| PANC-02-03 | AsPC-1 | CL1-0 | U266B1 |
|  |  |  | ACHN |

**Table S1.3:**
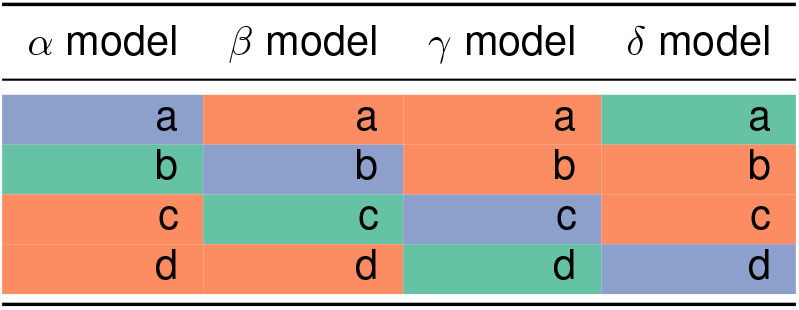
Four-fold cross-validation: Train and test sets for the four models and their respective holdout sets. The list of data sets for each split are summarized in table S1.2.

**Table S1.4:**
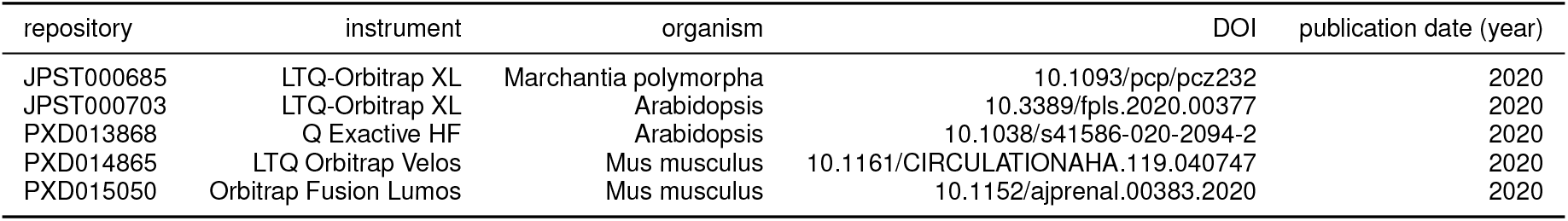
Details about validation data sets.

**Table S1.5:**
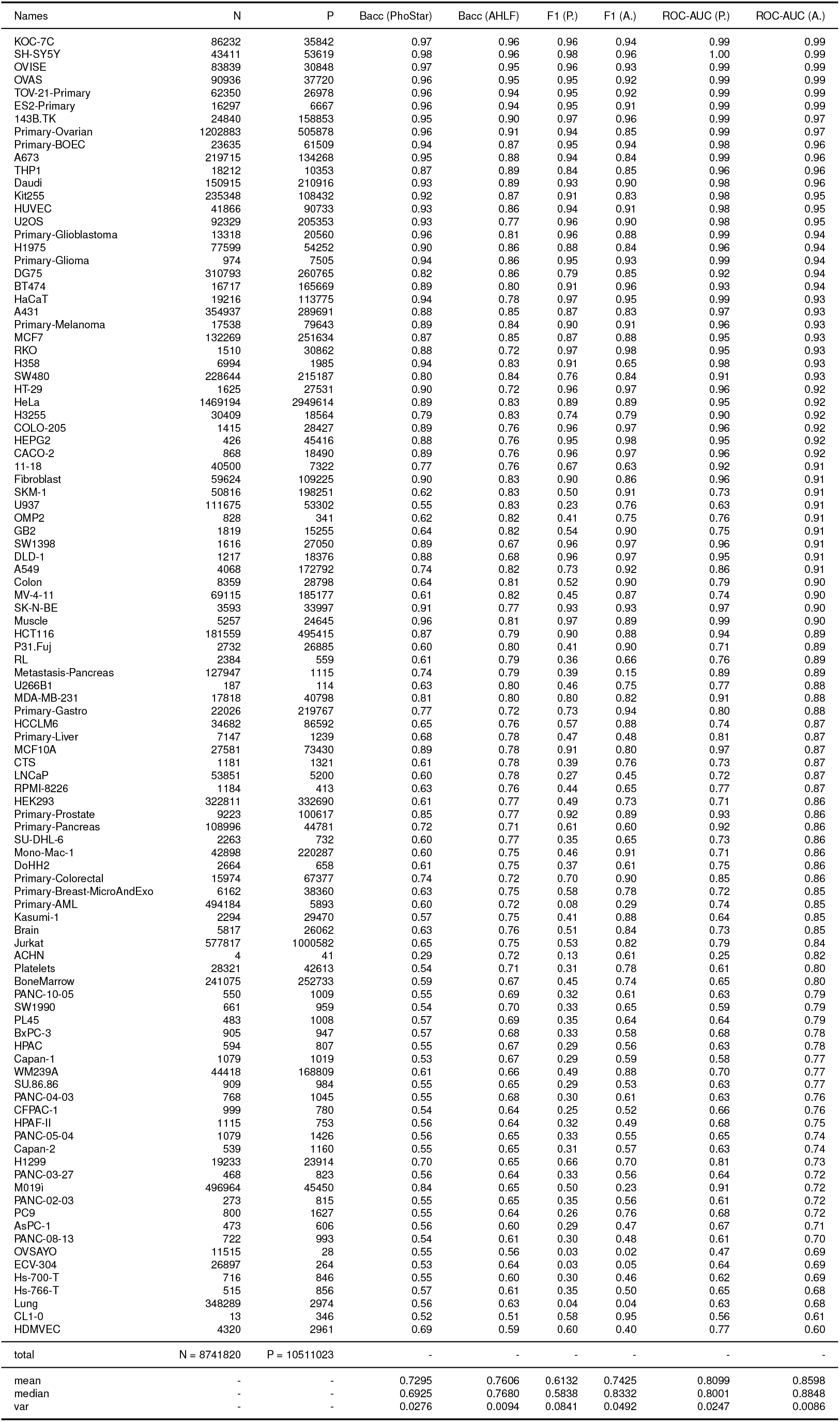
Metrics for the PXD012174 data set: balanced accuracy, F1 and ROC-AUC for PhoStar and AHLF. Rows are sorted according to ROC-AUC (AHLF) in descending order. For AHLF, metrics are computed on holdout-folds from cross-validation.

## S2 Supplementary Figures

**Figure S2.1:**
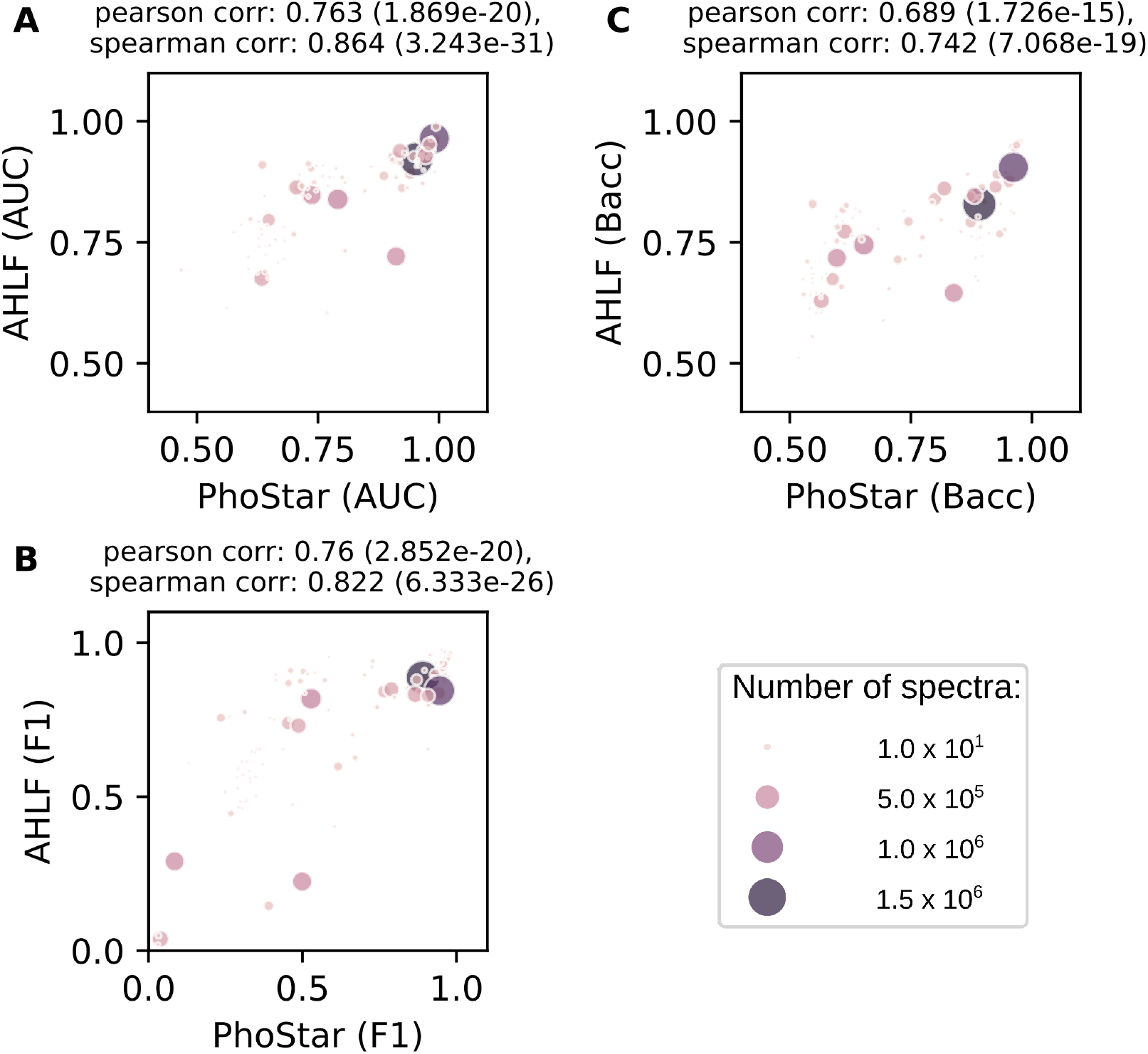
Correlation of performances between AHLF and PhoStar. **A-C:** ROC-AUC, F1-score and balanced accuracy of our method versus PhoStar. Each scatter plot is accompanied by Pearson and Spearman correlation coefficients (top) and p-values (top, in parentheses).

## S Model training and hyperparameter selection

Since this is the first end-to-end trained deep learning model directly applied to fragmentation spectra, no common architectures or even hyperparameters for this (or closely-related) problem set can be found from the literature. Hence, we report our experience with model training and particular choice of hyperparameters.

We trained our model on a V100 graphics cards from Nvidia. Each model took between 3-4 days to train. We trained on virtual epochs of 9000 steps. Each step is a gradient decent step on a mini-batch of size 64. A virtual-epoch contains a fraction of the complete training set (a true epoch contains all data points of the training set) and allows to evaluate the test error more frequently throughout the training process. From the trajectory of training-/test-loss over virtual epochs one can see that the model reaches acceptable values for binary accuracy ≈ 0.8 on the test samples after around 40 virtual-epochs. After these 40 virtual-epochs the training reaches a stable but slow reduction of training loss, while further improving the test set performance. We used dropout with a rate of 0.2 on the fully-connected layers of our model and could reduce overfitting significantly. In fact, one may continue training for more than 100 epochs and then start to see a test loss that stalls or increases implying overfitting. We used early stopping, but all (except one model) of our models trained for 100 virtual epochs, see supplemental Fig. S2.2. Considering regularization techniques, We only used dropout and early stopping as regularization. We chose an initial learning rate of 0.5·10^−6^ in combination with ADAM, an adaptive learning rate optimizer. We found the learning rate relatively sensitive. For example, a slightly smaller learning rate of 0.1·10^−6^ was ending up in an in-acceptable slow convergence, not shown here. In contrast, a slightly higher learning rate of 1.0·10^−6^ was showing not stable training due to high variance in training-loss and test-loss, not shown here. The number of layer in the block of dilated convolutions needs at least 8-10 layers, otherwise smaller models did not learn at all (training loss did not improve). We settled with 14 stacked-layers since additional layers lead to in-acceptable slow computation time for each backpropagation step on a mini-batch. A similar behavior could be seen for the number of filters. We settled with 64 filters per layer. With only 16 filters the the model was not able to learn. In contrast, with 128 filters per layers the time for training was in-acceptably slow. The latter hyperparameters (number of layers and filters) where tested for various learning rates in order to get an overview that is largely independent from the learning-rate. Of course this is one specific hyperparameter set out of many that may perform equally or even better. Still we hope that this summary of how we ended up with our particular parameter set may support future work.

**Figure S2.2:**
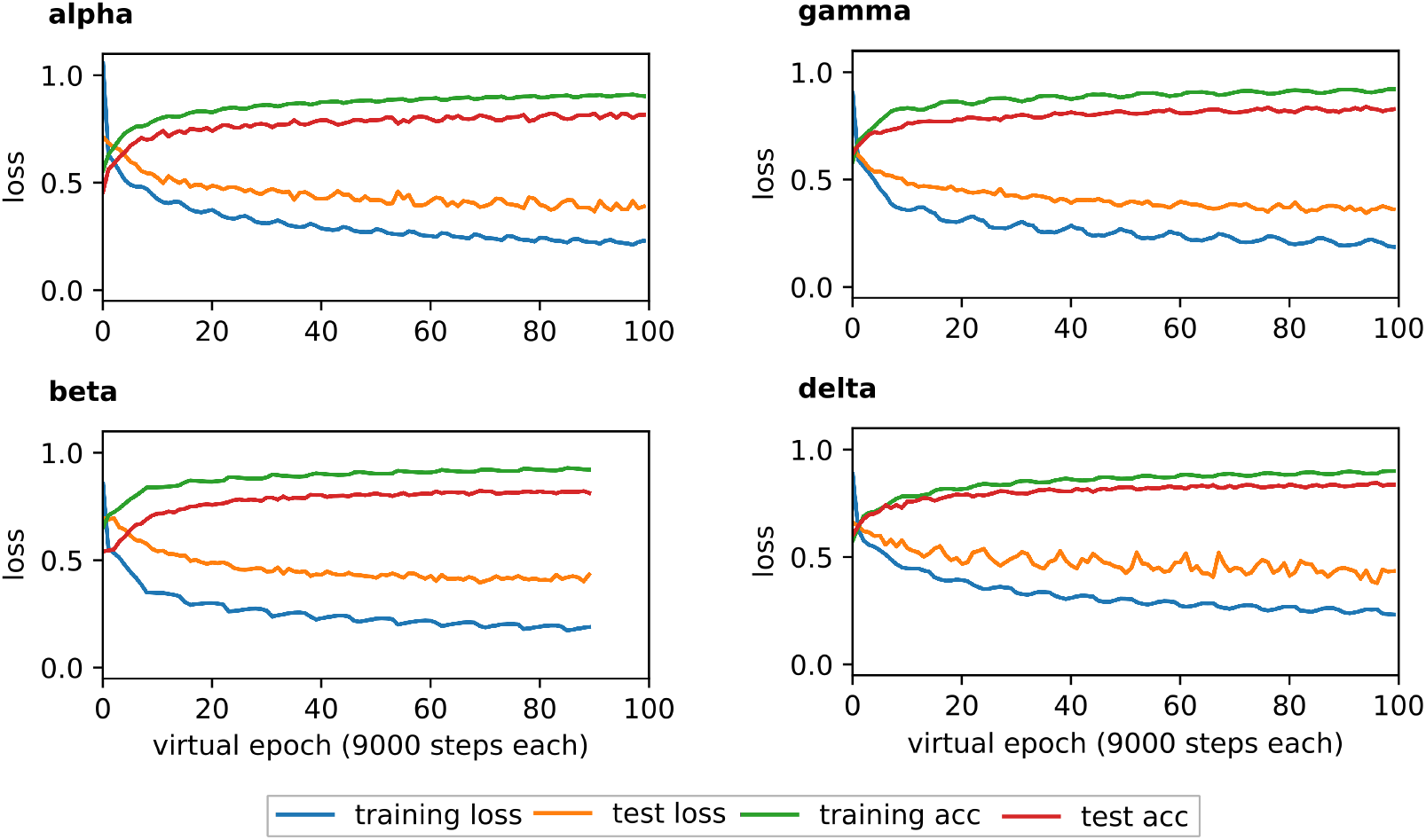
Training of four AHLF models due to four-fold cross-validation. **alpha-delta:** cross entropy loss for the models alpha, beta, gamma and delta. Each model was trained for 100 virtual epochs. Model beta stopped due to early stopping criteria at virtual epoch 89.

**Figure S2.3:**
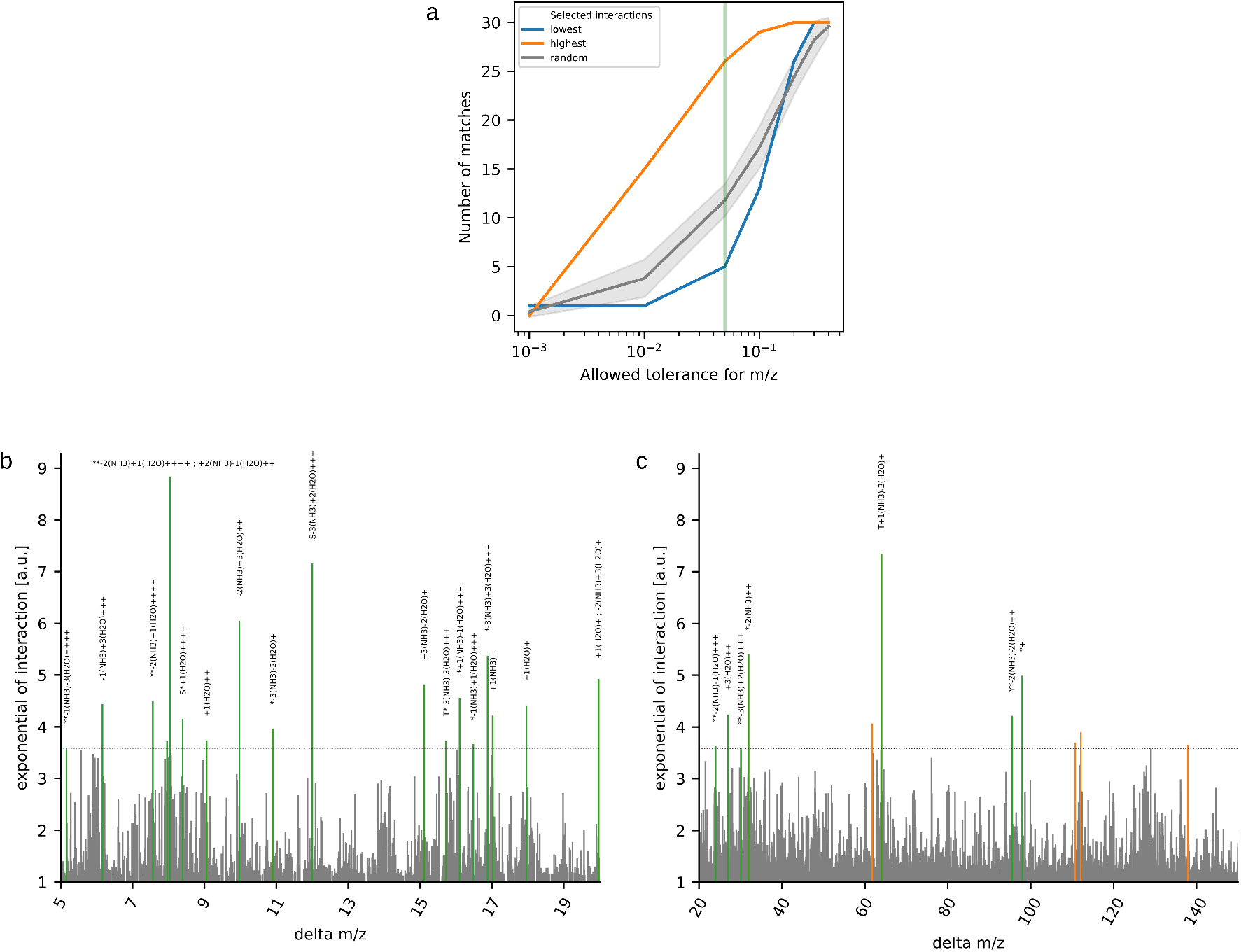
Matching delta m/z using a less strict tolerance. Similar to 4B-D from the main text, but instead here a higher tolerance was allowed when matching delta m/z. **a:** selecting tolerance of 0.05 m/z (green vertical line) as this yields more interactions in terms of absolute numbers while still relatively few potential false identifications (grey and blue line). **b+c:** 26 out of 30 highest interactions could be annotated (green peaks). Only the highest ten thousand interactions shown here.

**Figure S2.4:**
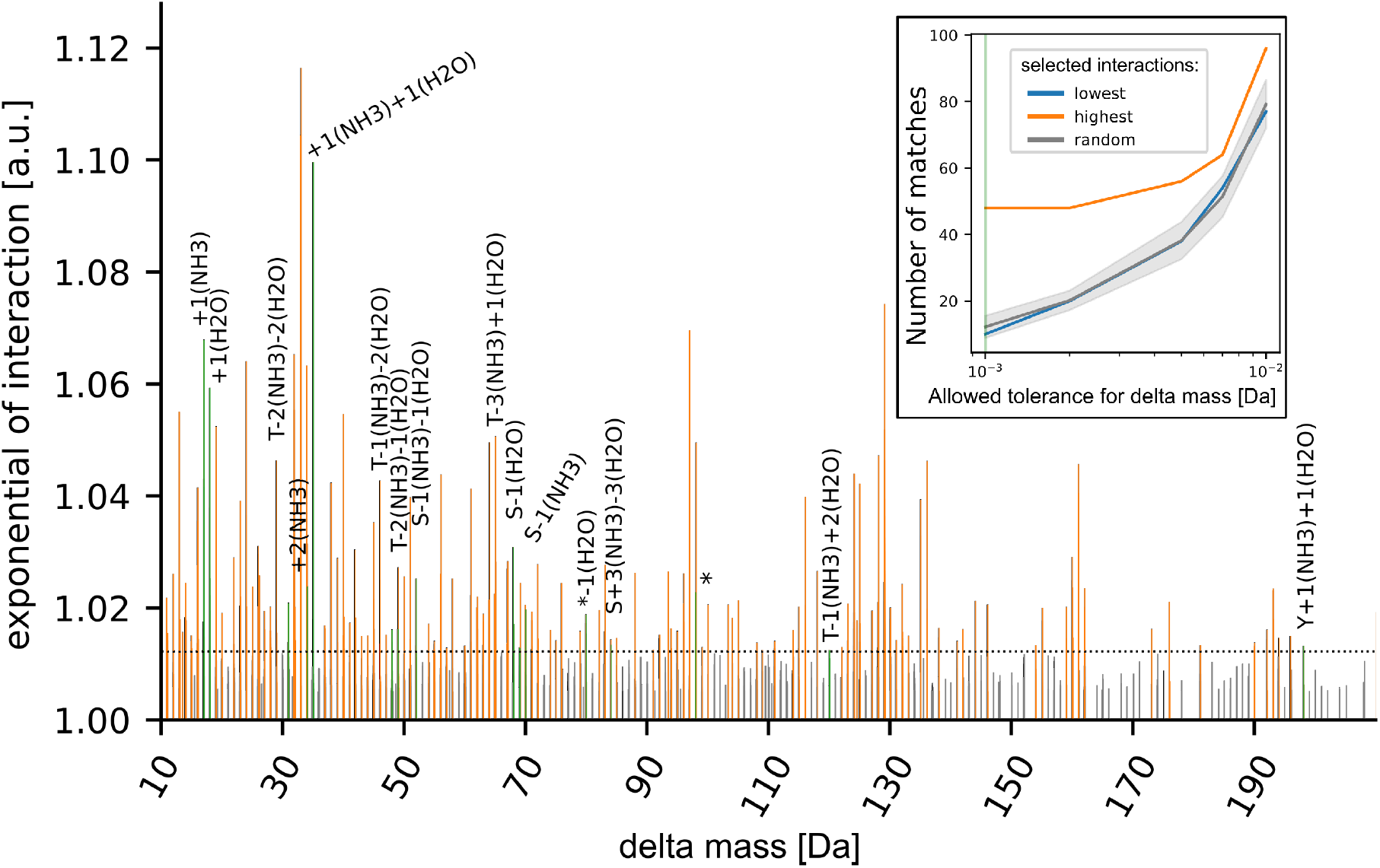
Matching delta masses for interactions for Orbitrap data for which spectra were deisotoped and de-charged. **Inlay:** selecting a tolerance of 0.001 Da (green vertical line) yields 48 matching interactions (with 16 unique delta masses) with few potential false identifications (grey and blue line). Sixteen unique delta masses out of top-500 highest interactions could be annotated (green peaks). Only the highest ten thousand interactions are shown here.

